# Igf signalling uncouples retina growth from body size by modulating progenitor cell division

**DOI:** 10.1101/2020.06.10.144360

**Authors:** Clara Becker, Katharina Lust, Joachim Wittbrodt

## Abstract

Balancing the relative growth of body and organs is of key importance for coordinating size and function. This is of particular relevance in post-embryonically growing organisms, facing this challenge life-long. We addressed this question in the neuroretina of medaka fish *(Oryzias latipes),* where growth and size regulation are crucial for functional homeostasis of the visual system. We find that a central growth regulator, Igf1 receptor, is necessary and sufficient for proliferation control in the postembryonic retinal stem cell niche, the ciliary marginal zone (CMZ). Targeted activation of Igf1r signalling in the CMZ uncouples neuroretina growth from body size control, increasing layer thickness while preserving the structural integrity of the retina. The retinal expansion is driven exclusively by enhanced proliferation of progenitor cells while stem cells do not respond to Igf1r modulation. Our findings position Igf signalling as key module controlling retinal size and structure with far reaching evolutionary implications.

## Introduction

During embryonic development and postembryonic life of multicellular organisms, the proportions of overall body and organ size are actively controlled, increased or decreased as necessary to sustain appropriate species-specific proportions. Scaling of overall body and organ size is known to be regulated by systemic signals, which couple nutritional status to growth (Andersen, Colombani, & Léopold, 2013). Conversely, differential growth of organs can be the result of altered sensitivity to systemic signalling (H. Y. Tang, Smith-Caldas, Driscoll, Salhadar, & Shingleton, 2011) as well as of variant intrinsic signalling in the respective organ (Bosch, Ziukaite, Alexandre, Basler, & Vincent, 2017; Twitty & Schwind, 1931).

To study organ size scaling, teleost fish such as medaka *(Oryzias latipes)* and zebrafish *(Danio rerio)* are particularly interesting model systems. They display life-long postembryonic growth and each organ harbours distinct stem cell populations, which continuously self-renew, generate progenitor and ultimately differentiated cells (Aghaallaei et al., 2016; Centanin et al., 2014; Furlan et al., 2017). In the eye and the retina precise regulation of continuous growth is of utmost importance to ensure the functional homeostasis of optical parameters and therefore vision. Differential eye size in teleost fish has been shown to be of functional relevance, since visual acuity is notably correlated with eye size (Caves, Sutton, & Johnsen, 2017). How retinal growth is regulated and uncoupled from body growth in different fish species to achieve differential eye sizes is currently not understood.

The teleost neuroretina contains a stem cell niche called the ciliary marginal zone (CMZ). The CMZ is located at the retinal margin and harbours the retinal stem and progenitor cells that continuously generate new neurons in teleosts and amphibians (Hollyfield, 1971; Johns & Easter, 1977; Raymond Johns, 1977; Straznicky & Gaze, 1971). Throughout postembryonic neurogenesis, multipotent, long-term self-renewing stem cells are located at the outermost periphery of the CMZ (Centanin et al., 2014; Raymond, Barthel, Bernardos, & Perkowski, 2006; Tsingos et al., 2019; Wan et al., 2016). Adjacent to retinal stem cells are rapidly dividing progenitor cells, which have restricted proliferative potential, are more heterogeneous and comprise several populations in different stages of lineage specification (Centanin et al., 2014; Pérez Saturnino, Lust, & Wittbrodt, 2018; Raymond et al., 2006; Wan et al., 2016).

A major integrator of organ size scaling within a growing organism is hormone signalling. Circulating hormones allow to translate extrinsic conditions such as nutrient availability into proportionate and coordinated growth of organs and body (Andersen et al., 2013; Boulan, Milán, & Léopold, 2015). The growth hormone signalling pathway centred around Insulin like growth factor (Igf) is well-known for its central role in regulating embryonic development and growth. A single nucleotide polymorphism in *igf1* present in small dog breeds was found to be a key factor for body size (Sutter et al., 2007). Mutations in *Igf1, Igf2* or *Igf1r* genes in mice, *ĩgflr* knockdown in zebrafish, as well as naturally occurring mutations in *Igf1* and *Igf1r* in humans lead to severe dwarfism phenotypes (Baker, Liu, Robertson, & Efstratiadis, 1993; Klammt, Pfäffle, Werner, & Kiess, 2008; Liu, Baker, Perkins, Robertson, & Efstratiadis, 1993; Schlueter, Peng, Westerfield, & Duan, 2007), underscoring the importance of growth hormone signalling in size determination. In insects, insulin-like peptides are important integrators of organ growth with developmental progression (Colombani, Andersen, & Léopold, 2012). Ultimately, final organ size is specified by systemic signals and organ-specific responses, which are defined by discrete insulin sensitivity of each organ (Shingleton & Frankino, 2018). The expression and localisation of Igf pathway components during teleost embryonic and larval development has been assessed in a variety of species (Ayaso, Nolan, & Byrnes, 2002; Boucher & Hitchcock, 1998; Radaelli et al., 2003a; Radaelli, Patruno, Maccatrozzo, & Funkenstein, 2003b; Zygar, Colbert, Yang, & Fernald, 2005). Consistently, Igf ligands and receptors (Igfr) were found to be expressed in the retina in developmental and adult stages. The most detailed analysis stems from studies in postembryonic goldfish retinae where *igf1* is expressed in the retina and its binding sites are localised in the CMZ and IPL (Boucher & Hitchcock, 1998; Otteson, Cirenza, & Hitchcock, 2002). In zebrafish, Igf1r-mediated signalling is required for proper embryonic development, especially of anterior neural structures, and inhibition results in reduced body size, growth arrest and developmental retardation (Eivers, McCarthy, Glynn, Nolan, & Byrnes, 2004; Schlueter et al., 2007). Additionally, Igf1r knockdown leads to increased neuronal apoptosis, reduced proliferation and adversely affects cell cycle progression (Schlueter et al., 2007).

The role of Igf signalling in retinal stem cells has been studied in the postembryonic retina of chicken and quails where Igf1 or insulin injection increases proliferation in the CMZ (Fischer & Reh, 2000; Kubota, Hokoc, Moshiri, McGuire, & Reh, 2002). Despite the expression of its pathway components in the CMZ, the role of Igf1r signalling in the teleost CMZ as well as the consequences of its alteration for the continuously growing neuroretina have not been addressed.

In this study, we elucidate the function of Igf1r signalling in the postembryonic retinal stem cell niche of the medaka fish. We establish that consistent with other fish species, signalling pathway components are expressed in the retina and specifically the CMZ. We find that global Igf inhibition reduces proliferation in the CMZ. Targeted, constitutive activation of Igf1r signalling specifically in CMZ stem and/or progenitor cells increases proliferation and leads to a prominent increase in eye size and uncouples eye size from body size. Importantly, the enlarged retinae are structurally intact and properly laminated, indicating that progenitor differentiation and neurogenesis is not disturbed. We demonstrate that Igf1r activation decreases cell cycle length in CMZ cells, expands the progenitor population and ultimately leads to increased neuronal cell numbers. By dissecting the individual contribution of stem cells and progenitor cells to eye size increase we uncover that only retinal progenitor but not stem cells are responsive to modulation by Igf1r signalling.

## Results

### Igf1r signalling regulates proliferation in the medaka CMZ

The retinal stem cell niche is located in a continuous ring in the CMZ at the periphery of the teleost eye (Fig. 1A). To address the involvement of the Igf signalling pathway in regulating retinal stem and progenitor cell proliferation in the medaka retina, we first assessed expression of receptors and ligands. We generated probes for several genes, of which *igf1ra, igf2* and *insrb* were reliably expressed in the CMZ as well as in specific layers of the differentiated retina (Fig. S1).

**Fig. 1:**
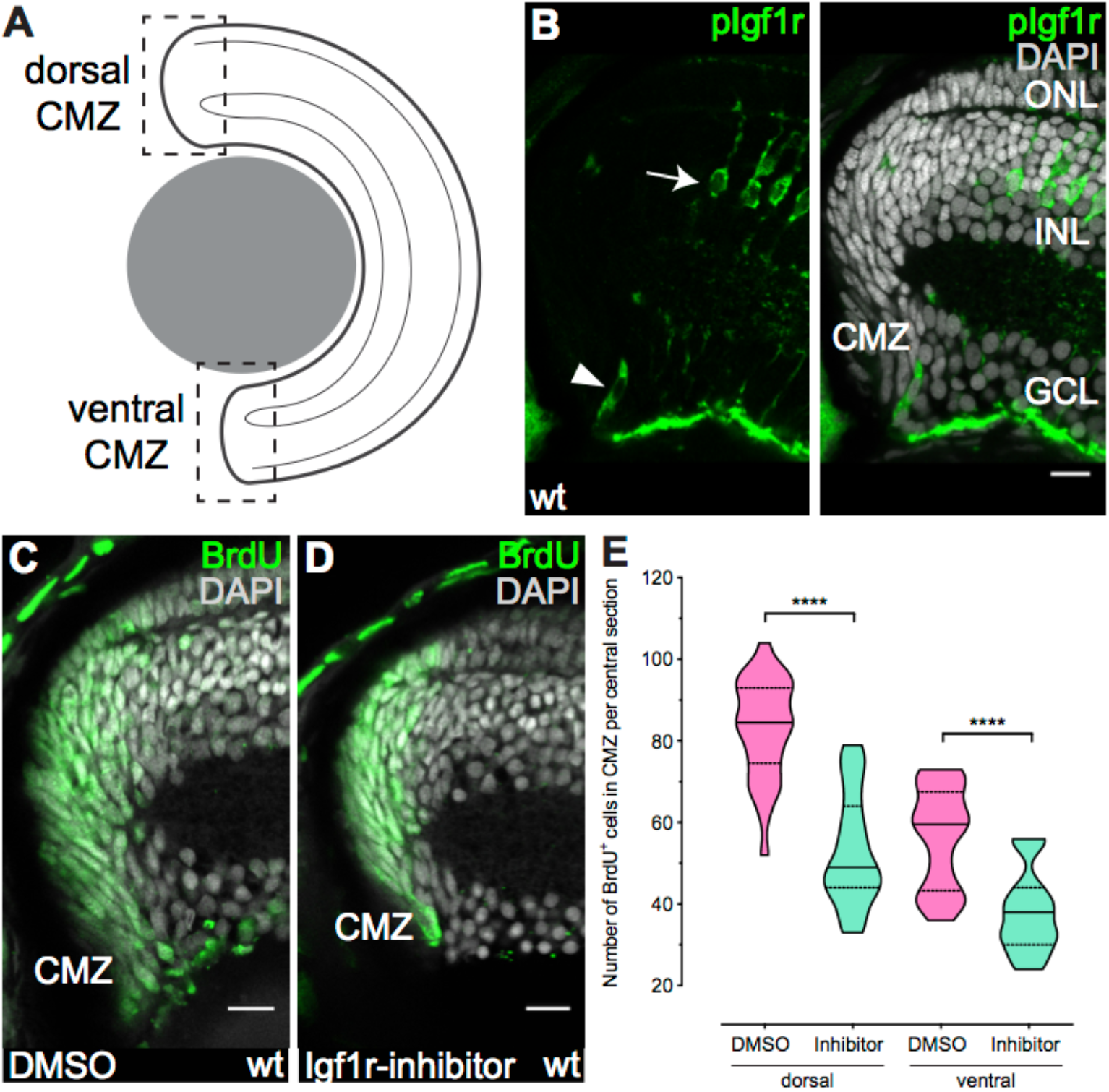
Igf1r signalling regulates proliferation in the CMZ. (A) Schematic representation of a retinal section. Dashed squares represent dorsal (d) and ventral (v) CMZ. All CMZ sections in this paper depict the dorsal CMZ, with quantifications being done separately for dorsal (CMZ_d_) and ventral (CMZ_v_) CMZ due to inherent differences in proliferation, marker expression and morphology. (B) Cryosection of wildtype (wt) hatchling with anti-pIgf1r (green) staining shows that Igf1r is active in single cells in the CMZ (arrowhead, n = 65 cells in 58 sections from 10 retinae) and in Müller glia cells (arrow) in the inner nuclear layer (INL). Scale bar is 10 μm; outer nuclear layer (ONL). (C-D) Wt hatchlings were incubated for 24 h in BrdU and 10 μM Igf1r inhibitor NVP-AEW541 or DMSO. Cryosections of DMSO- (C) and Igf1r-inhibitor-treated (D) retinae with BrdU staining (green) display decreased BrdU incorporation upon Igf1r inhibition. Scale bars are 10 μm. (E) Quantification of the number of BrdU-positive cells in one Z plane per central section shows a decrease in Igf1r-inhibitor-treated retinae (n = 26 (dorsal)/23 (ventral) sections from 10 retinae) compared to DMSO (n = 28 (dorsal)/27 (ventral) sections from 10 retinae) (median + quartiles, ****P_d/v_ < 0.0001).

The activity of receptor tyrosine kinases like Igf1r is mediated by ligand-dependent dimerisation and subsequent trans-phosphorylation. To assess whether Igf1r is active in the CMZ, we performed immunostainings against phosphorylated Igf1r (pIgf1r). Single cells in the progenitor domain of the CMZ as well as Müller glia cells in the inner nuclear layer (INL) in the differentiated part of the retina were positive for pIgf1r (Fig. 1B).

Based on our Igf1r expression and activity data, we hypothesised that Igf signalling is involved in regulating proliferation in the CMZ. To determine the impact of Igf1r signalling on retinal stem and progenitor cell proliferation, we made use of the widely-used Igf1r inhibitor NVP-AEW541 (Chablais & Jazwinska, 2010; W.-Y. Choi et al., 2013; Huang et al., 2013) and BrdU to label S phase cells. Fish at hatching stage were incubated in 10 μM NVP-AEW541 or DMSO together with BrdU for 24 h, and analysed afterwards by immunostaining against BrdU (Fig. 1C, D). Inhibition of Igf1r signalling resulted in a 30 % decrease of S phase cells in the CMZ (Fig. 1E), validating that Igf1r-mediated signalling is crucial for proliferation in the CMZ. These results show that ligands and receptors of the Igf signalling cascade are expressed and active in the CMZ and that Igf signalling activity is necessary for CMZ proliferation.

### Constitutive activation of Igf1r in the CMZ results in increased eye size

Based on the previous results, we hypothesized that Igf1r signalling represents a promising target to modulate differential growth of the retina. We therefore sought to induce precise continuous activation of Igf1r signalling in the CMZ.

We generated a transgenic line where a constitutively active *cd8a:igf1ra* chimeric receptor (*caigf1r*) is expressed under the control of the *rx2* promoter, a retina-specific transcription factor. The *cd8a:igf1ra* variant was generated by an in-frame fusion of the extracellular and transmembrane domain of medaka *cd8a* and the intracellular domain of medaka *igf1ra*, as previously described (Carboni et al., 2005). The *rx2* promoter drives expression in stem cells and early multipotent progenitors in the CMZ as well as in Müller glia and photoreceptor cells in the differentiated retina (Reinhardt et al., 2015).

We validated the ability of the *rx2::caigf1r* transgenic line to activate the signalling cascade downstream of Igf1r in *rx2*-positive cells by examining the phosphorylation status of Akt, a known downstream component of IGF signalling. We therefore performed immunostainings against pAkt on cryosections. In retinae of wildtype hatchlings, pAkt was present in the peripheral domain in the CMZ (Fig. S2A). In *rx2::caigf1r* retinae pAkt staining covered a larger area and appeared brighter, overlapping with the *caigf1r* expression domain (Fig. S2B). Strong pAkt signal was also evident in photoreceptor cells, which express *caigf1r* as well. These results establish that the *rx2::caigf1r* transgenic line is able to activate Akt downstream of Igf1r in the medaka CMZ.

To address the potential of Igf1r signalling to induce differential retinal growth we next examined hatchlings for changes in retinal morphology related to *rx2::caigf1r* expression. Intriguingly, transgenic *rx2::caigf1r* hatchlings displayed a prominent increase in eye size compared to wildtype siblings (Fig. 2A-C), with otherwise normal head size and body length (Fig. S3A-C). Relative eye size (anterior-posterior eye diameter normalised to body length) was significantly increased in *rx2::caigf1r* hatchlings (Fig. 2D). The increase in relative eye size persisted throughout postembryonic growth until adulthood (Fig. 2E).

**Fig. 2:**
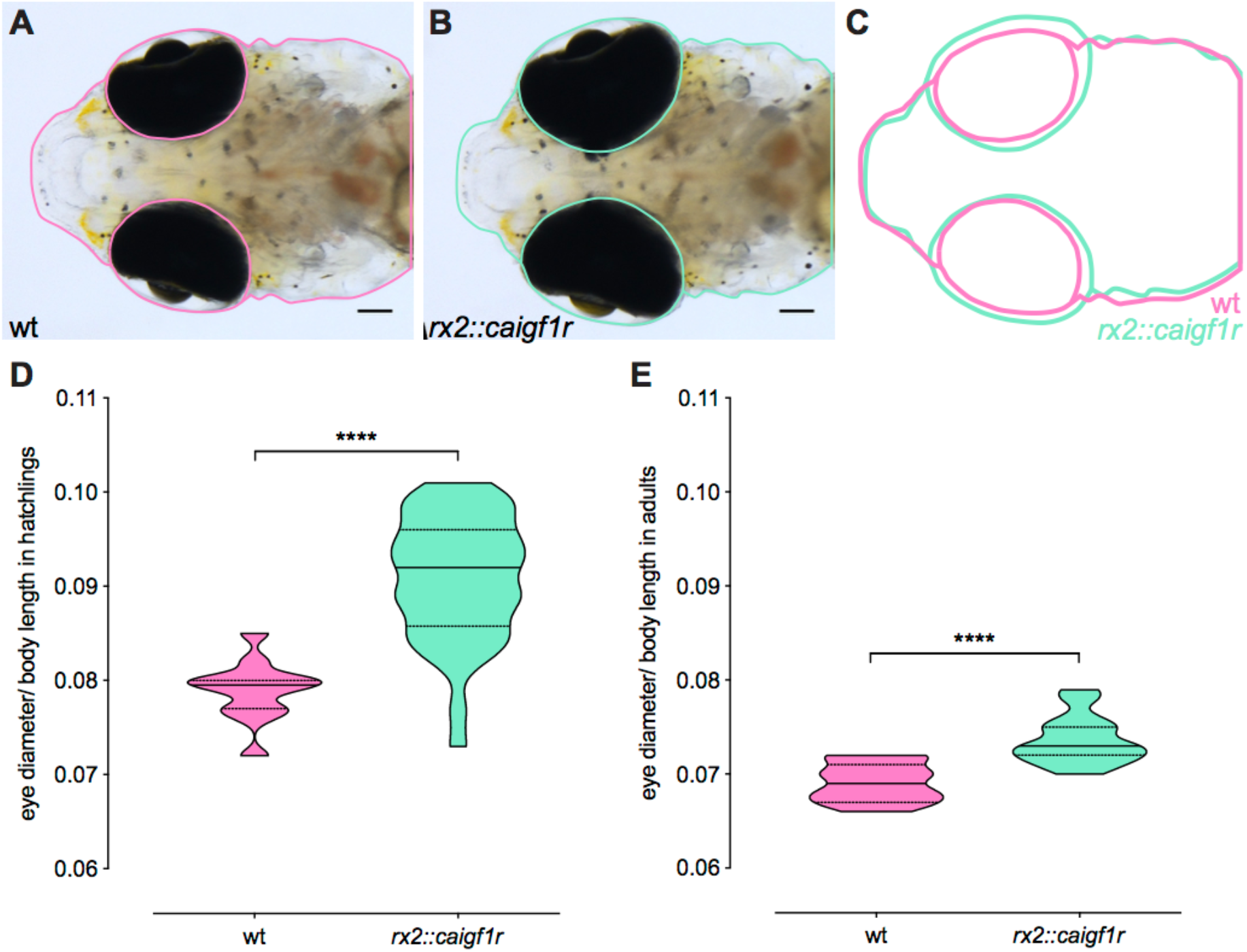
Constant activation of Igf1r in retinal stem and progenitor cells results in increased eye size. (A-C) Eye size of *rx2::caigf1r* hatchlings (B) is larger compared to wt siblings (A). Scale bars are 100 μm. (D-E) Quantification of relative eye size (eye diameter normalised to body length) of wt (D: n = 12; E: n = 11) and *rx2::caigf1r* (D: n = 38; E: n = 15) hatchlings (D) and adults (E) (median + quartiles, ****P < 0.0001).

This data shows that CMZ-targeted activation of Igf1r signalling is able to uncouple retinal growth from overall body growth in medaka.

### Retinal enlargement stems from neuroretinal expansion through increase in cell number

Size increase of a tissue can arise due to different mechanisms of tissue expansion, such as increase in cell size or number, or stretching and increase in fluid or pressure (Ritchey, Zelinka, Tang, Liu, & Fischer, 2012; Stujenske, Dowling, & Emran, 2011; Veth et al., 2011). To understand how Igf1r signalling could mediate eye size increase, we examined cryosections of wildtype and *rx2::caigf1r* hatchling retinae. *Rx2::caigf1r* eyes exhibited a prominent expansion of the neuroretina in contrast to wildtype controls (Fig. 3A, B). Importantly, nuclear morphologies and arrangement indicated that the overall retinal architecture in *rx2::caigf1r* fish remained intact. The stereotypical structure of the neuroretina with the CMZ at the periphery and three nuclear and two plexiform layers in the differentiated part was undisturbed by retinal expansion.

**Fig. 3:**
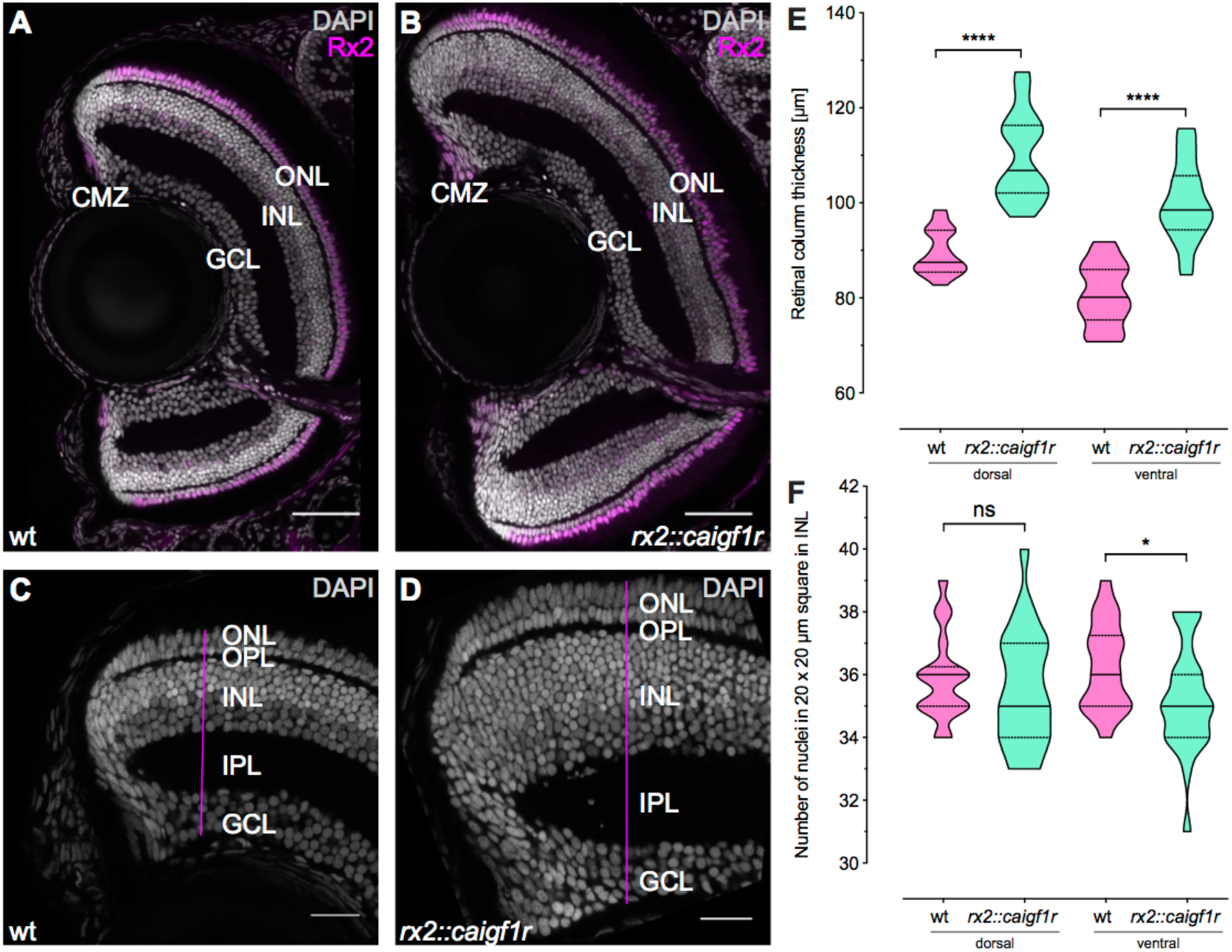
Retinal enlargement stems from neuroretinal expansion through increase in cell number. (A-B) Cryosections of wt (A) and *rx2::caigf1r* (B) hatchling retinae with staining against Rx2 (magenta) display neuroretinal expansion. Scale bars are 50 μm. (C-D) Thickness measurements were done along a line (magenta) perpendicular to the inner plexiform layer (IPL). Thickness of the whole retinal column and all individual layers were measured in the fully laminated part close to the CMZ in wt (n = 18 sections from 12 retinae) and *rx2::caigf1r* (n = 24 sections from 14 retinae) retinae. Scale bars are 20 μm. (E) Quantification of retinal column thickness in the dorsal as well as ventral retina shows increase in *rx2::caigf1r* (n = 24 sections from 14 retinae) compared to wt (n = 18 sections from 12 retinae) retinae (median + quartiles, ****P_d/v_ < 0.0001). (F) Quantification of nucleus number in a 20 x 20 μm INL region shows similar amounts in *rx2::caigf1r* (n = 24 sections from 14 retinae) and wt (n = 18 sections from 12 retinae) retinae (median + quartiles, ^ns^P_d_ = 0.5031, *P_V_ = 0.0456).

We assessed retinal topology further using the expression of *rx2*. Both peripheral *rx2* expression in the CMZ as well as central expression in Müller glia cells and photoreceptors showed an identical pattern between wildtype and *rx2::caigf1r* retinae, indicating that these cell populations are present (Fig. 3A, B). To characterise neuroretina expansion in more detail, we measured the thickness of the neuroretina in *rx2::caigf1r* and wildtype retinae. Measurements of the retinal column were taken in the peripheral but fully laminated region of the retina, along a line perpendicular to the inner plexiform layer (IPL) (Fig. 3C, D). Retinal column thickness was increased by 20 μm on average in the dorsal as well as ventral retina (Fig. 3E). We additionally measured retinal column thickness in central regions, which reflect the embryonic contribution as *rx2* is also expressed in retinal progenitor cells during development. Thickness of the central neuroretina in *rx2::caigf1r* hatchlings was also increased compared to wildtype. However, the increase was greatest in more peripheral CMZ-derived regions, arguing for a greater impact of the postembryonic contribution through the CMZ (Fig. S4A). Analysis of the thickness of individual neuroretina layers showed that all nuclear layers and the outer plexiform layer (OPL) increased their thickness. The most prominent expansion took place in the INL (Fig. S4B-F), accounting for up to 15 μm of the 20 μm increase in *rx2::caigf1r* retinae. In contrast, the IPL showed decreased thickness in the ventral retina (Fig. S4C). To determine whether the expansion was due to enlarged cell size, we quantified the number of nuclei in a 20 x 20 μm square region in the INL as an approximation for cell size. The number of nuclei remained constant between wildtype and *rx2::caigf1r* retinae (Fig. 3F), indicating that cell size is not enlarged.

Taken together, these results show that the activation of Igf1r signalling in the CMZ increases retinal size, and the enlargement stems from neuroretinal expansion through an increase in cell number rather than cell size.

### Igf1r signalling activation decreases cell cycle length in the CMZ

Igf1r signalling has been shown to influence cell cycle progression in different *in vitro* and *in vivo* models (Hodge, D’Ercole, & O’Kusky, 2004; Schlueter et al., 2007). To determine whether Igf1r signalling activation in *rx2::caigf1r* fish affects the cell cycle of proliferating cells in the CMZ, we next analysed cell cycle and S phase length of the retinal progenitor population in wildtype and *rx2::caigf1r* hatchlings. To this end, we deployed an established experimental regime using BrdU and EdU pulses (Das, Choi, Sicinski, & Levine, 2009; Klimova & Kozmik, 2014). Hatchlings were incubated in BrdU for 2 h, washed, and incubated in EdU for 30 min before analysis. Immunostainings against BrdU, EdU and Pcna (Fig. 4A, B) allowed to quantify different fractions of single and double-positive cells, from which cell cycle and S phase length were calculated. In *rx2::caigf1r* retinae, the cell cycle length was decreased from 12 h to 10.5 h on average (Fig. 4C), whereas S phase length remained constant with an average duration of 4.5 h (Fig. 4D). Moreover, in *rx2::caigf1r* compared to wildtype fish, the number of BrdU-positive cells in the CMZ was more than doubled (Fig. 4E).

**Fig. 4:**
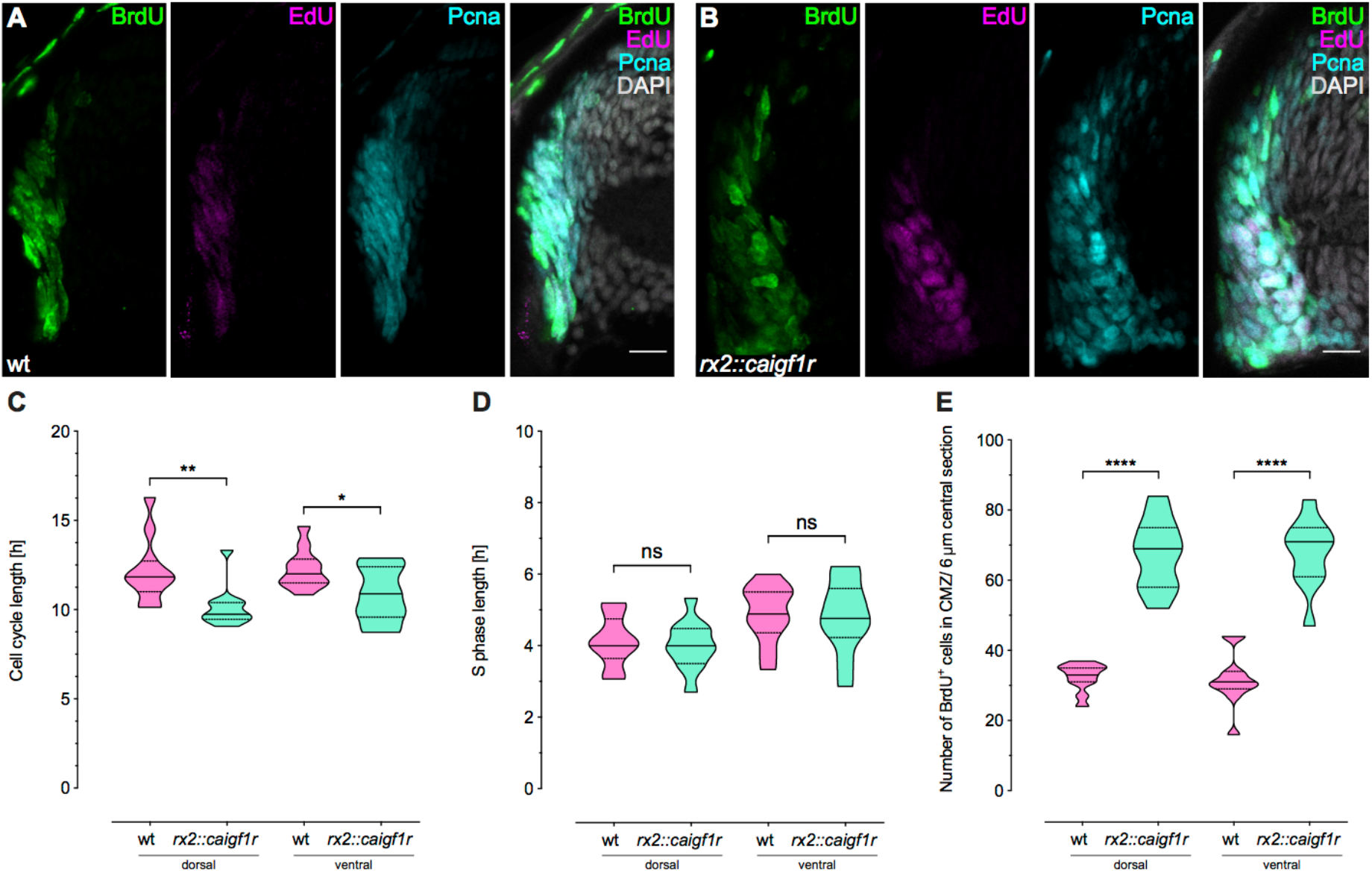
Constant activation of Igf1r signalling decreases cell cycle length in the CMZ. (A-B) Cryosections of wt (A) and *rx2::caigf1r* (B) hatchling retinae incubated for 2 h in BrdU and 30 min in EdU to determine cell cycle length. Staining against BrdU (green), EdU (magenta) and Pcna (cyan) show partial overlap in the CMZ. Scale bars are 10 μm. (C) Quantification of cell cycle length shows a reduction of 1-2 h in *rx2::caigf1r* (n = 11 sections from 4 retinae) compared to wt (n = 11 sections from 4 retinae) retinae (median + quartiles, **P_d_ = 0.0053, *P_v_ = 0.0188). (D) Quantification of S phase length in *rx2::caigf1r* (n = 11 sections from 4 retinae) compared to wt (n = 11 sections from 4 retinae) retinae (median + quartiles, ^ns^P_d_ = 0.6764, ^ns^P_v_ = 0.8223). S phase length is not altered in *rx2::caigf1r* retinae. (E) Quantification of BrdU-positive cell number in the CMZ per 6 μm central section shows that numbers have more than doubled in *rx2::caigf1r* (n = 11 sections from 4 retinae) compared to wt (n = 11 sections from 4 retinae) retinae (median + quartiles, ****P_d/v_ < 0.0001).

Taken together, these data show that in *rx2::caigf1r* hatchlings, more cells in the CMZ go through the cell cycle faster, thereby increasing retinal cell numbers resulting in increased retinal size and expanded retinal layers.

### Activation of Igf1r signalling in the CMZ increases retinal progenitor but not stem cell number

We observed a relative eye size increase when specifically activating Igf signalling both in stem and progenitor cells by expressing the constitutively activated receptor under the control of the stem and progenitor cell-specific promoter *rx2 (rx2::caigf1r).* Differential responses of retinal stem and progenitor cells to extrinsic stimuli have been previously described (Centanin et al., 2014; Love, Keshavan, Lewis, Harris, & Agathocleous, 2014), which prompted us to disentangle the contribution of each cell population to the expansion in response to Igf1r signalling modulation. Hence, we sought a more specific driver to target IGF modulation exclusively to stem cells. We therefore analysed the cytosolic non-specific dipeptidase *zgc:114181* (hereafter *cndp*) promoter, which was found to have promising CMZ-restricted retinal expression but was not characterised in detail (Haas, Wittbrodt and Wittbrodt, unpublished).

We characterised the 5 kb promoter region upstream of *cndp* by generating transgenic reporter lines *(cndp::eGFP-caax, cndp::H2A-mCherry). Cndp* expression was detected in the retina (Fig. 5A) and the choroid plexi in the brain (Fig. S5A, B). In the retina, *cndp*-positive cells were found exclusively in a small, peripheral subset of Rx2-positive cells in the CMZ (Fig. 5A). To functionally validate the potential of *cndp-expressing* cells, we employed a Cre/loxP-mediated lineage tracing approach. We generated a construct where a Tamoxifen-inducible Cre^ERT2^ was expressed under the control of the *cndp* promoter (Fig. 5B) and combined it with the GaudíRSG red-switch-green reporter line (Centanin et al., 2014). GaudíRSG embryos were injected with the *cndp::Cre^ERT2^* plasmid at 1-cell stage and induced with Tamoxifen at hatchling stage. After 2-3 weeks, fish were analysed for GFP expression by whole-mount immunostaining (Fig. 5C). Retinae displayed GFP-positive clones that originated in the CMZ and were continuous to the differentiated retina (Fig. 5D), while also extending into the ciliary epithelium. These clones labelled cells in all three nuclear layers (Fig. 5E) and thus were categorised as <u>i</u>nduced <u>A</u>rched <u>C</u>ontinuous <u>S</u>tripes (iArCoS, (Centanin et al., 2014)). Importantly, we did not observe clonal footprints originating from progenitor cells which establishes cndp as a bona fide marker for neuroretinal stem cells.

**Fig. 5:**
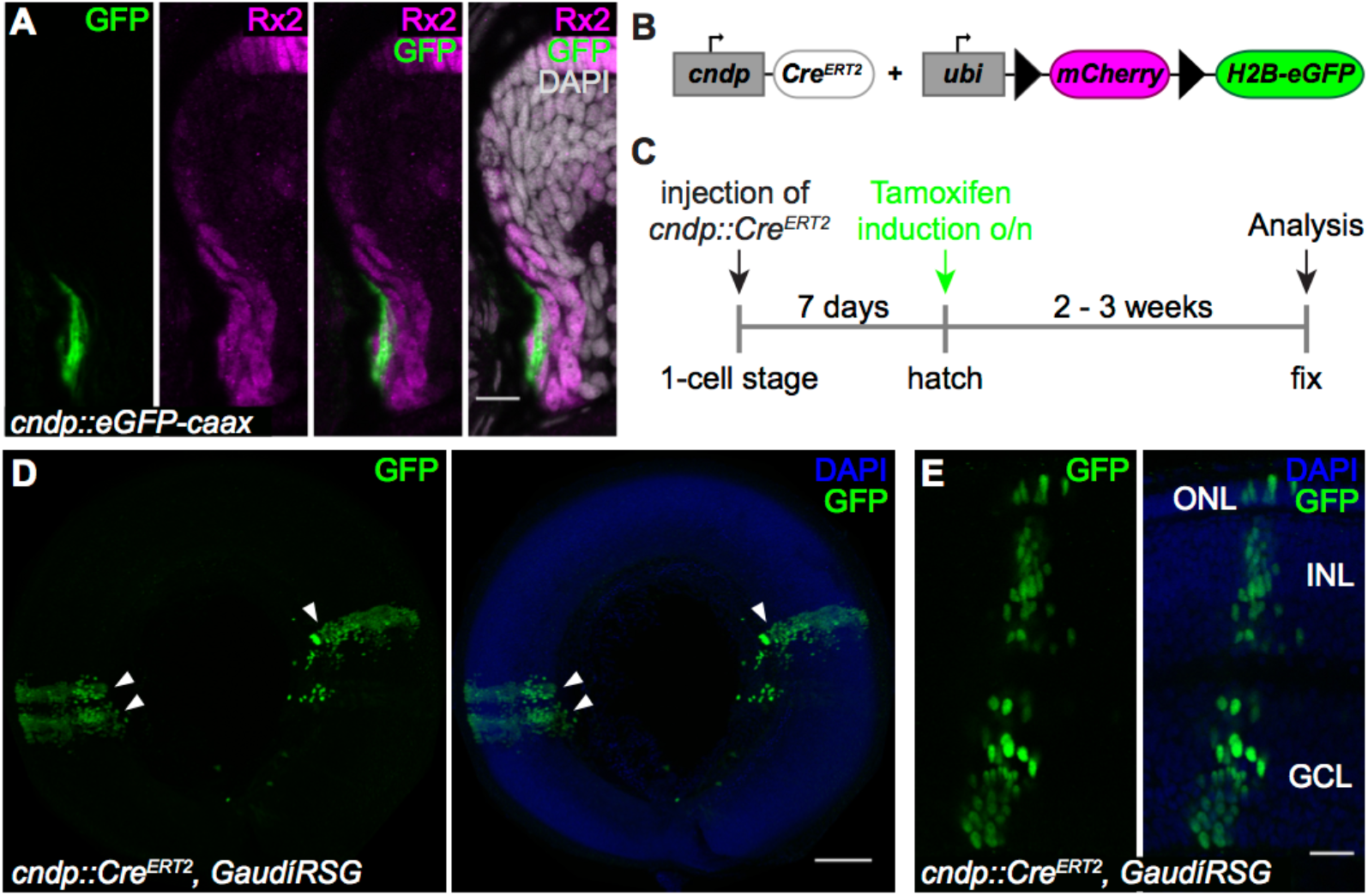
*Cndp* is expressed in multipotent neuroretinal stem cells. (A) Cryosection of a *cndp::eGFP-caax* hatchling retina with staining against GFP (green) in a peripheral subset of the Rx2 (magenta) domain in the CMZ. Scale bar is 10 μm. (B) Schematic representation of the constructs used for lineage tracing. Upon tamoxifen induction, *mCherry* is floxed out and *H2B-eGFP* is expressed in GaudíRSG fish. (C) Experimental outline: *cndp::Cre^ERT2^* is injected in 1-cell stage GaudíRSG embryos. At hatch, fish are incubated in tamoxifen overnight and grown for 2 – 3 weeks before analysis. (D-E) Whole-mount immunostainings of *cndp::Cre^ERT2^*, GaudíRSG retinae against GFP (green) with neuroretinal clones (D) labelling the whole retinal column (E) and extending into the ciliary epithelium, originating from multipotent neuroretinal stem cells (arrowheads) (n = 7 clones in 3 retinae). Scale bars are 100 μm (D) and 20 μm (E).

To understand whether stem and progenitor cells possess similar responsiveness to activated Igf1r signalling we made use of the specific expression domains of *cndp* and *rx2* in different retinal stem *(cndp, rx2)* and progenitor (*rx2*) cell populations (Fig. 6A). The *cndp::H2A-mCherry* reporter line was used to identify retinal stem cells, and the Rx2 antibody to label both retinal stem and progenitor cells (Reinhardt et al., 2015).

**Fig. 6:**
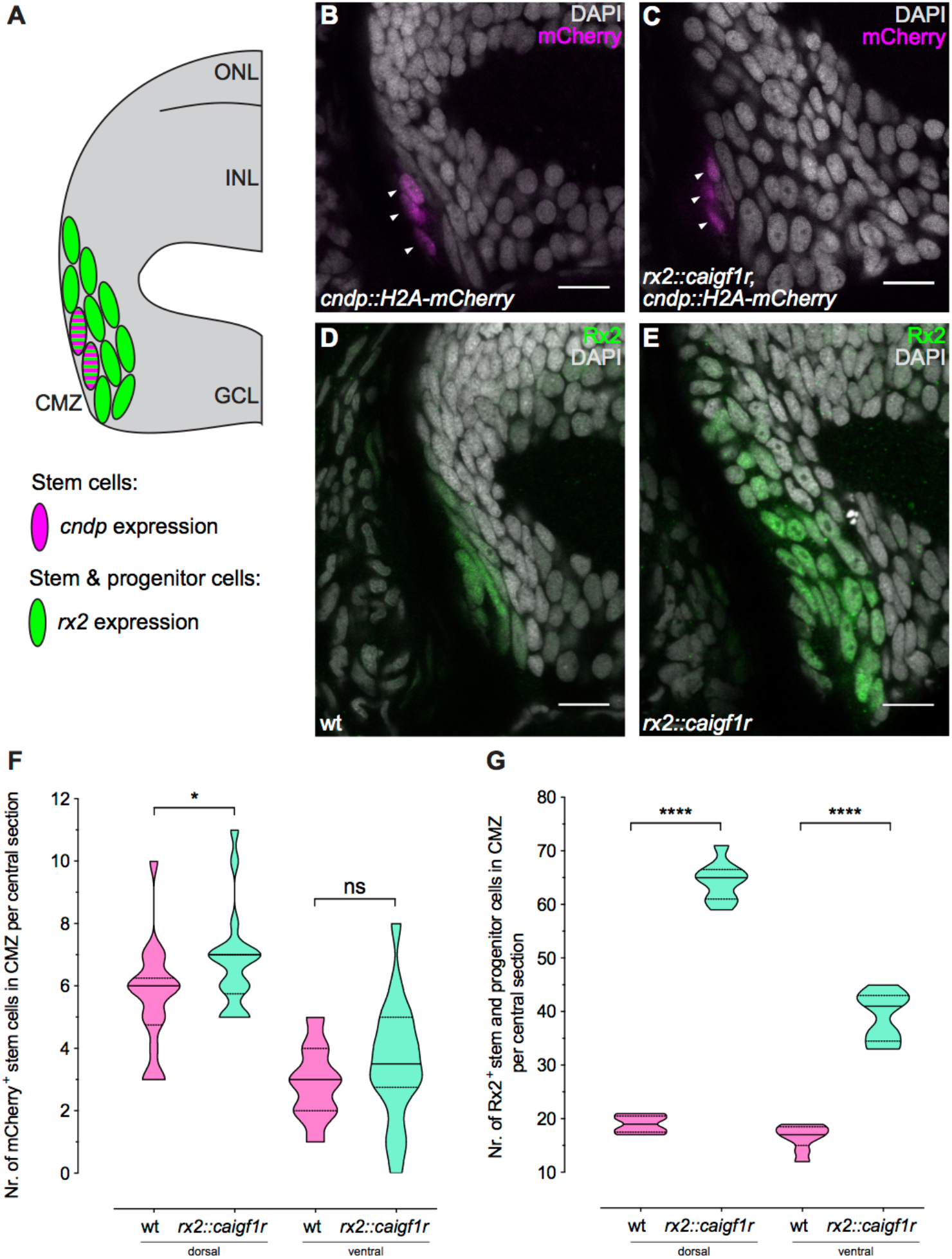
Constant activation of Igf1r signalling expands retinal progenitor cell numbers. (A) Schematic representation of the dorsal CMZ of a retinal section with *cndp* (magenta) and *rx2* (green) expression in stem and progenitor cells. (B-C) Cryosections of wt (B) and *rx2::caigf1r* (C) *cndp::H2A-mCherry* reporter hatchling retinae. mCherry (magenta) is visible in peripheral-most cells in the CMZ (arrowheads). (D-E) Cryosections of wt (D) and *rx2::caigf1r* (E) hatchling retinae. Rx2 staining (green) marks peripheral cells in the CMZ. (F) Quantification of H2A-mCherry-positive cell number in the CMZ of 16 μm central sections does not indicate an expansion of *cndp*-positive stem cells in *rx2::caigf1r* (n = 18 sections from 6 retinae) compared to wt (n = 18 sections from 6 retinae) retinae (median + quartiles, *P_d_ = 0.0442, ^ns^P_v_ = 0.2177). (G) Quantification of Rx2-positive cell number in the CMZ of 16 μm central sections demonstrates that Rx2-positive stem and progenitor cells have more than doubled in *rx2::caigf1r* (n = 9 sections from 6 retinae) compared to wt (n = 9 sections from 6 retinae) retinae (median + quartiles, ****P_d/v_ < 0.0001).

We assessed the number of retinal stem cells expressing *cndp* by immunostainings on wildtype and *rx2::caigf1r* hatchlings, positive for the *cndp::H2A-mCherry* reporter (Fig. 6B, C). In both wildtype and *rx2::caigf1r* retinae the number of *cndp*-positive stem cells per section was low, ranging from 3 to 11 in the dorsal and 0 to 8 in the ventral CMZ. The number of *cndp*-positive stem cells was rather stable, with a slight increase in *rx2::caigf1r* retinae (Fig. 6F). In contrast to that, the Rx2 domain was prominently expanded in *rx2::caigf1r* versus wildtype hatchlings (Fig. 6D, E), with Rx2-positive stem and progenitor cells in the CMZ being more than doubled in *rx2::caigf1r* retinae (Fig. 6G), arguing that the progenitor, but not the stem cell population is expanded by Igf1r signalling activation.

Since we found almost no increase in stem cell numbers in retinae of *rx2::caigf1r* hatchlings, we next wanted to understand whether retinal stem cells are not responding to Igf1r signalling modulation. We generated a transgenic line in which Igf1r signalling activation is targeted to *cndp*-positive retinal stem cells and expressed *caigf1r* under the control of the stem cellspecific *cndp-promoter (cndp::caigf1r).* Functionality of the construct was evident through the enlargement of the choroid plexi compared to wildtype siblings (Fig. S6A, B). However, the GFP expression domain in the retina was unaltered (Fig. S6C). Additionally, we examined relative eye size in hatchlings of two independent lines derived from different founders. Neither *cndp::caigf1r* line displayed an alteration in relative eye size compared to its wildtype siblings (Fig. S6D). These results demonstrate that the *cndp*-expressing retinal stem cell population does not expand upon Igf1r signalling activation. This further confirms and refines our findings that only the progenitor but not stem cell population is expanded in *rx2::caigf1r* retinae.

Taken together, we demonstrate that retinal growth can be uncoupled from overall body growth through the activation of Igf1r signalling targeted to the CMZ. This intrinsic modulation elicits differential responses in stem and progenitor cell populations in the medaka retina, leading to a shortened cell cycle and consequential increase of retinal progenitor but not stem cell numbers.

## Discussion

In this study, we investigated how the growth of an organ can be uncoupled from overall body growth in medaka fish. We focused on the retina and dissected the role of Igf1r signalling in regulating proliferation of the retinal stem cell niche and changes in retina size. We found that the retinal stem cell niche is permissive for mitogenic signalling mediated by Igf1r activity. Using a combination of expression analysis and gain-as well as loss-of-function approaches, we examined the function of Igf1r signalling in the postembryonic retinal stem cell niche of medaka. Ligands and receptors of the Igf pathway are expressed in the postembryonic retina and specifically in the CMZ, indicating a local paracrine signalling hub. Furthermore, the pathway is active in sparse progenitors in the CMZ. Inhibition of Igf1r signalling leads to decreased proliferation of retinal stem and progenitor cells. CMZ-targeted constitutive activation results in a dramatic increase of eye size originating from cell number increase. We determined that this effect is caused by the specific expansion of the progenitor cell pool by speeding up the cell cycle without affecting the subsequent differentiation potential, while retinal stem cells do not respond to Igf1r signalling activation (Fig. 7).

**Fig. 7:**
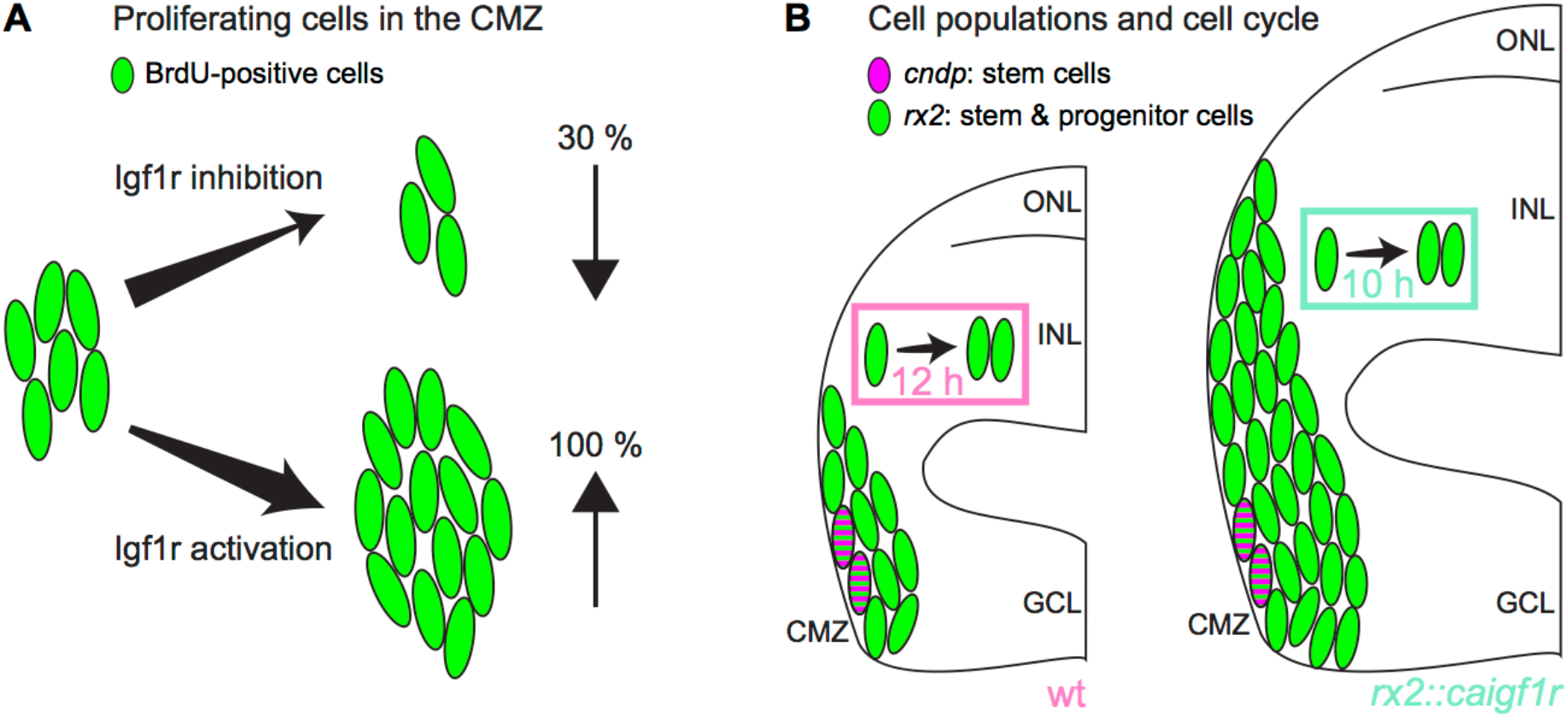
Igf1r signalling regulates proliferation and progenitor population size in the CMZ. (A) Igf1r inhibition decreases proliferating cells (green) in the CMZ by 30 %, while Igf1r activation increases proliferation by ≥ 100 %. (B) Igf1r activation in the CMZ in *rx2::caigf1r* fish expands progenitor numbers (*rx2*-positive, green), but does not enlarge the stem cell population (*cndp*-positive, magenta). Cell cycle in the CMZ is shortened from 12 h in wildtype fish to 10 h upon Igf1r activation in *rx2::caigf1r* fish.

We showed that constitutive activation of Igf1r signalling expands only the *rx2*-positive progenitor but not stem cell population. Targeting stem cells by specific Igf1r activation in *cndp*-expressing cells does not impact on retinal size, indicating that retinal stem cells do not respond to Igf1r signalling with increased proliferation. In contrast, progenitor numbers are more than doubled in response to constant activation of Igf1r signalling, rendering the *rx2*-positive progenitor population receptive for this mitogenic stimulus. Since slowly dividing stem cells act as long-lasting reservoir for a continuous and secure replenishment of rapidly cycling progenitor cells, their requirements differ with regard to ensuring genomic stability and integrity in order to prevent whole lineages from acquiring detrimental mutations. This difference could be explained by distinctive downstream signal transduction and absence thereof, or specific safeguarding of stem cells against unwarranted mitogenic stimuli. Tumour suppressor genes are likely candidates to impede deviant proliferation. For example, adult *p53* knockout mice display increased neural stem cell proliferation in the subventricular zone, supporting a regulatory role of p53 in controlling proliferation in this niche (Gil-Perotin et al., 2006; Meletis et al., 2006).

Differential responses of stem and progenitor cells have been observed after nutrient deprivation in the *Xenopus* retina. Whereas stem cells are resistant to nutrient deprivation and mTOR inhibition, retinal progenitors respond with changes in proliferation and differentiation in an mTOR-mediated manner (Love et al., 2014). These differences between retinal stem and progenitor cells are similar to our observations where stem cells are not responsive to a mitogenic stimulus mediated by Igf1r, while progenitor proliferation is increased. It is important to note that we observe a local, cell-intrinsic effect of a targeted stimulus, while Love and colleagues examine a global effect on a local cell population. In the future, it will be exciting to profile transcriptional changes in retinal stem cells in our model of Igf1r activation to address whether stem cells resist to changes by adaptation of intrinsic signalling networks or whether they are unable to be changed at all.

Upon Igf1r signalling activation, the thickness of the neuroretina drastically increases. Intriguingly, enlarged retinae are structurally intact displaying proper lamination and differentiation. While retinal enlargement due to enhanced progenitor proliferation might be an expected phenotype, we were surprised by the perfect morphological arrangement of retinal cells in *rx2::caigf1r* fish. Interestingly, upon *Yap* mRNA injection in *Xenopus* embryos, tadpoles display enlarged eyes with more proliferating cells in the retina, but retinal morphology and lamination is disturbed (Cabochette et al., 2015). Conversely, Yap knockdown leads to a decrease in S phase length, whereas the overall cell cycle length is increased. We observed that Igf1r expression leads to overall cell cycle shortening, but the S phase is not affected, which is in agreement with results from various systems where Igf signalling influences G1 or G2 phase length (Hodge et al., 2004; Schlueter et al., 2007; S. Wang, Wang, Wu, & Han, 2015). Increase in retinal size has also been observed in a zebrafish retina mutant for *patched2* where retinal patterning and retinal morphology are largely intact, only Müller glia numbers are negatively affected (Bibliowicz & Gross, 2009). Furthermore, retinal progenitor cells in the CMZ are expanded while keeping a constant cell cycle length. Contrastingly, we observed a shortened cell cycle in the progenitor population which is likely causal for its expansion in *rx2::caigf1r* fish.

Teleost species display a great variety in retinal size, architecture and cell type composition dependent on their photic environment and habitat. Surface-dwelling fish like medaka have two layers of photoreceptors, one light-sensitive rod and one cone layer responsible for colour vision, while zebrafish possess one layer of cones and already three to four layers of rods for enhanced light perception, as they live in deeper waters (Lust & Wittbrodt, 2018). Interestingly, retinae of many deep-sea fish possess predominantly rods at the expense of other retinal neurons, resulting in high rod to ganglion cell convergence for enhanced light sensitivity (Darwish, Mohalal, Helal, & El-Sayyad, 2015; de Busserolles, Fitzpatrick, Marshall, & Collin, 2014; Wagner, Fröhlich, Negishi, & Collin, 1998). Whether the functionality of the enlarged retina in *rx2::caigf1r* fish concerning circuitry, visual acuity and light sensitivity is the same as in wildtype fish remains to be studied. We did not observe apparent behavioural changes in these fish, however testing for example responsiveness to visual stimuli as well as visual acuity measurements could give insights into adaptations resulting from eye size change.

The isolated size increase of an apparently functional sensory organ leads us to speculate about the evolutionary and ecological significance of retinal size and architecture. Throughout the teleost clade, different adaptations to specific habitats and niches are evident in the retina. One particularly interesting example is the four-eyed fish *Anableps anableps*, which displays structural differences within the retina to accommodate its specific optic requirements. This species features eyes that partly protrude over the top of their skull and are above the water surface while the lower half of the eye is submerged just below the surface. Subsequently, composition and thickness of the retina differs between ventral and dorsal halves, which are equipped for aerial and aquatic vision, respectively. The ventral retina features an INL that is twice as thick as the dorsal INL, concordant with increased proliferation in the ventral compared to the dorsal CMZ during larval development (Perez et al., 2017). Therefore, proliferative regulation and relative cell type composition in the differentiated retina must differ in ventral and dorsal halves. As we found a pronounced increase in INL thickness upon Igf1r signalling activation, it is tempting to speculate that Igf signalling plays a role in manifesting the structural differences in the *Anableps anableps* retina. In the teleost retina, several populations of lineage-specified progenitors reside in the CMZ, and modification of their transcriptional signatures shift cell type ratios (Pérez Saturnino et al., 2018). Based on the preferential accumulation of INL cells in the ventral retina of *Anableps anableps*, one possible scenario is that a progenitor population lineage-committed to generate INL cells is expanded and proliferates more.

The susceptibility of retinal progenitor cells to altered Igf signalling might permanently modify retinal architecture, ultimately facilitating the rapid occupation of new ecological niches followed by subsequent speciation. We propose that Igf signalling can act as an evolutionary hub, through which retinal size, morphology and cell type composition is altered by modifying signalling activity in distinct populations of progenitor cells in the CMZ.

## Materials and Methods

### Animals and transgenic lines

Medaka (*Oryzias latipes*) used in this study were kept as closed stocks at Heidelberg University. All experimental procedures and husbandry were performed in accordance with the German animal welfare law and approved by the local government (Tierschutzgesetz §11, Abs. 1, Nr. 1, husbandry permit AZ 35–9185.64/BH and line generation permit AZ 35–9185.81/G-145-15). Fish were maintained in a constant recirculating system at 28°C on a 14 h light/10 h dark cycle. The following stocks and transgenic lines were used: wildtype Cabs, *Heino* mutants (Loosli et al., 2000)*, rx2::caigf1r rx2::lifeact-eGFP, cndp::H2A-mCherry, cndp::eGFP-caax, cndp::H2B-eGFP, cndp::caigf1r cndp::H2B-eGFP, cndp::Cre^ERT2^, GaudíRSG* (Centanin et al., 2014). All transgenic lines were created by microinjection with Meganuclease (I-SceI) in medaka embryos at the one-cell stage, as previously described (Thermes et al., 2002).

The constitutively active Igf1r variant (caIgf1r, Cd8a:Igf1ra) was generated by an in-frame fusion of the codon-optimised extracellular and transmembrane domain of olCd8a (synthesised by Geneart) and the intracellular domain of olIgf1ra, as previously described (Carboni et al., 2005). The *cndp::Cre^ERT2^* plasmid was generated by cloning the 5 kb *cndp* regulatory region in a pBS/I-SceI-vector containing a tamoxifen-inducible Cre recombinase. The plasmid contains *cmlc2::eCFP* as insertional reporter.

### BrdU/EdU incorporation

For BrdU incorporation, hatchlings were incubated in 2.5 mM BrdU (Sigma-Aldrich) diluted in 1x embryo rearing medium (ERM, 17 mM NaCl, 40 mM KCl, 0.27 mM CaCl_2_, 0.66 mM MgSO_4_, 17 mM Hepes) for 2 h. For EdU incorporation, hatchlings were incubated in 250 μM EdU (ThermoFisher) diluted in 1x ERM for 30 min. Quantification of BrdU-positive cells was performed in four retinae from individual hatchlings. Cell counts were performed in z = 6 μm of two to three central sections per retina.

### Igf1r inhibition

For inhibition of Igf1r, hatchling fish were incubated in 10 μM NVP-AEW541 (Selleckchem) diluted in 1x ERM at 28°C for 24 h. In a parallel control group, hatchling fish were incubated in 0.001 % DMSO/1x ERM at 28°C for 24 h. Directly afterwards fish were euthanised and fixed for analysis.

### Induction of Cre/lox system

For Cre^ERT2^ induction, hatchlings were treated with a 5 μM tamoxifen solution (Sigma-Aldrich) in 1x ERM overnight.

### Immunohistochemistry on cryosections

Fish were euthanised using 20x Tricaine and fixed overnight in 4 % PFA, 1x PTW at 4°C. After fixation samples were washed with 1x PTW and cryoprotected in 30 % sucrose in 1x PTW at 4°C. To improve section quality, the sections were incubated in a half/half mixture of 30 % sucrose and Tissue Freezing Medium for at least 3 days at 4°C. 16 μm thick serial sections were obtained on a cryostat. Sections were rehydrated in 1x PTW for 30 min at room temperature. Blocking was performed for 1-2 h with 10 % NGS (normal goat serum) in 1x PTW at room temperature. The respective primary antibodies were applied diluted in 1 % NGS o/n at 4°C. The secondary antibody was applied in 1 % NGS together with DAPI (Sigma-Aldrich, D9564; 1:500 dilution in 1x PTW of 5 mg/ml stock) for 2-3 h at 37°C. Slides were mounted with 60 % glycerol and kept at 4°C until imaging.

### BrdU and Pcna immunohistochemistry on cryosections

BrdU and Pcna antibody staining was performed with an antigen retrieval step. After all antibody stainings and DAPI staining, except for BrdU/Pcna, were complete, a fixation for 30 min was performed with 4 % PFA. Slides were incubated for 1.5 h at 37°C in 2 N HCl solution, and pH was recovered by washing with a 40 % Borax solution in 1x PTW before incubation with the primary BrdU or Pcna antibody.

### EdU staining on cryosections

EdU staining reaction was performed after all other antibody stainings were completed using the Click-iT™ EdU Alexa Fluor™ 647 Flow Cytometry Assay Kit according to manufacturer’s protocol (Thermo Fisher).

### Immunohistochemistry on wholemount retinae

Fish were euthanised using 20x Tricaine and fixed overnight in 4 % PFA in 1x PTW at 4°C. After fixation, samples were washed with 1x PTW. Fish were bleached with 3 % H_2_O_2_, 0.5 % KOH in 1x PTW for 2-3 h in the dark. Retinae were enucleated and permeabilised with acetone for 15 min at −20°C. Blocking was performed in 1 % bovine serum albumin (Sigma-Aldrich), 1 % DMSO (Roth/Merck), 4 % sheep serum (Sigma-Aldrich) in 1x PTW for 2 h. Samples were incubated with primary antibody in blocking buffer overnight at 4°C. The secondary antibody was applied together with DAPI in blocking buffer overnight at 4°C. Primary antibodies were used at 1:200, secondary antibodies at 1:250 and DAPI at 1:500.

### Antibodies

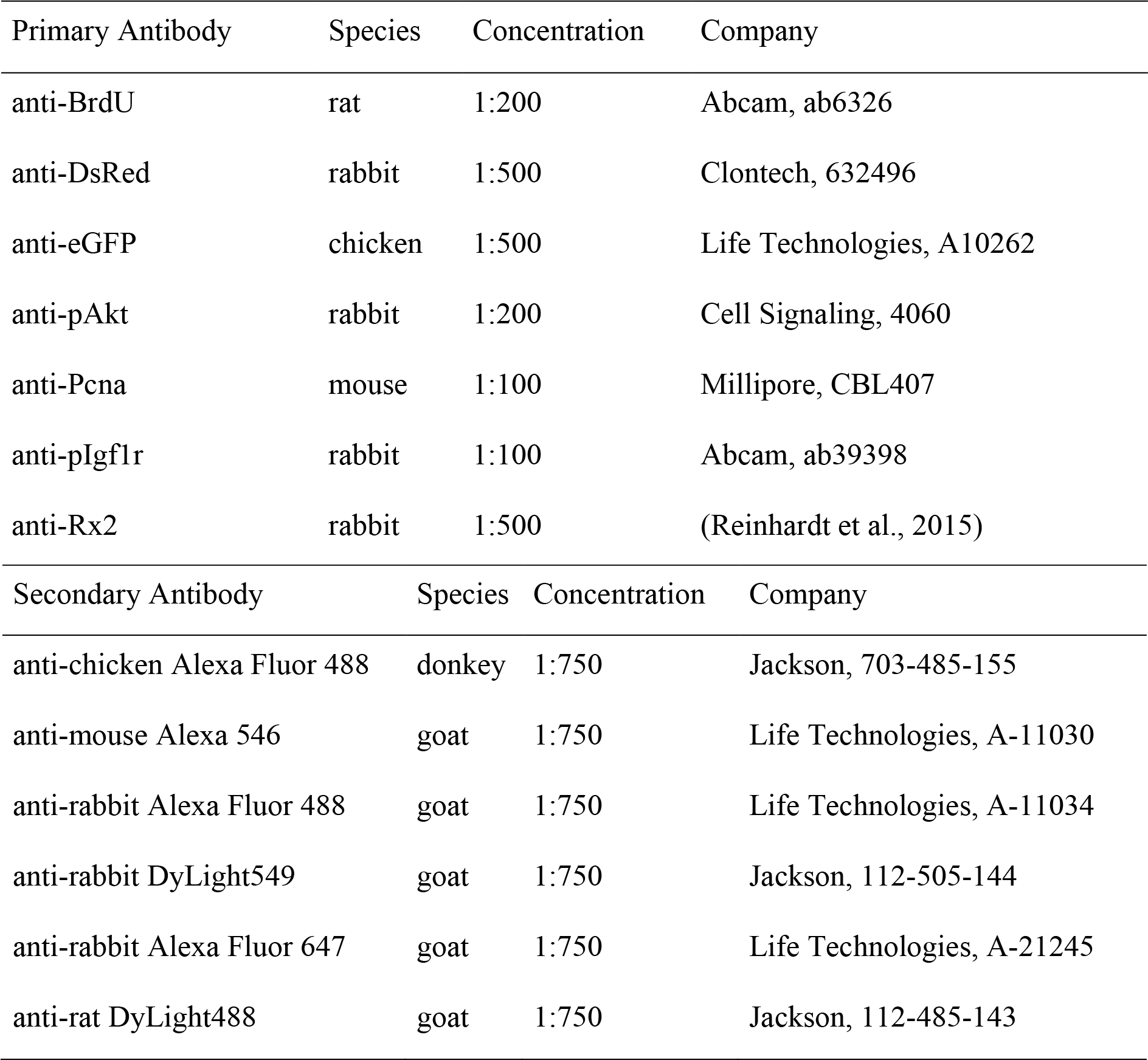

### Measurement of cell cycle and S phase length

To determine cell cycle length of retinal progenitor cells, BrdU and EdU were used as previously described (Das et al., 2009). Hatchlings were incubated for 2 h in BrdU, then 30 min in EdU before fixation. Pcna antibody staining was used to label all cycling retinal progenitor cells. Pcna-, EdU- and BrdU-positive cells as well as cells positive for only BrdU were quantified. Cell cycle length and S phase length were determined: T_cell cycle_ = 2 h *(Pcna^+^ cells/BrdU^+^ only cells); T_S phase_ = 2 h *(EdU^+^ cells/BrdU^+^ only cells). Four retinae from individual hatchlings were used for analysis. Cell counts were performed in z = 6 μm of two to three central sections per retina. T_cell cycle_ and T_S phase_ were determined for individual sections.

### Wholemount *in situ* hybridisation

Wholemount *in situ* hybridisations using NBT/BCIP detection were carried out as previously described (Loosli, Köster, Carl, Krone, & Wittbrodt, 1998). Afterwards, samples were cryoprotected in 30 % sucrose in 1x PTW overnight at 4°C and 20 μm thick serial sections were obtained on a Leica cryostat. Sections were rehydrated in 1x PTW for 30 min at room temperature and washed several times with 1x PTW. Slides were mounted with 60 % glycerol and kept at 4°C until imaging.

### Image acquisition

All immunohistochemistry images were acquired by confocal microscopy at a Leica TCS SP8 with 20x or 63x glycerol objective. Sections of wholemount *in situ* hybridisations were imaged at a Zeiss Axio Imager M1 microscope. Images of whole hatchlings were acquired with a Nikon SMZ18 Stereomicroscope equipped with the camera Nikon DS-Ri1.

### Image processing and statistical analysis

Images were processed via Fiji image processing software. Statistical analysis and graphical representation of the data were performed using the Prism software package (GraphPad). Violin plots show median, 25th and 75th percentiles. Unpaired two-tailed t-tests were performed to determine the statistical significances. The P value P < 0.05 was considered significant and P values are given in the figure legends. Sample size (n) is mentioned in every figure legend. No statistical methods were used to predetermine sample sizes, but our sample sizes are similar to those generally used in the field. The experimental groups were allocated randomly, and no blinding was done during allocation.

## Acknowledgements

We thank the Wittbrodt lab for constructive discussions on the project, Alex Cornean, Steffen Lemke, Tinatini Tavhelidse, Erika Tsingos, Venera Weinhardt and Lucie Zilova for critical reading of the manuscript. We are grateful to A. Saraceno, E. Leist and M. Majewski for fish husbandry. C.B. and K.L. were members of HBIGS, the Heidelberg Graduate School for Life Sciences. C.B. was supported by a PhD fellowship of the Studienstiftung des deutschen Volkes. This work was supported by the European Research Council (GA 294354-ManISteC to J.W.) and the Deutsche Forschungsgemeinschaft (SFB 873 TP A3 to J.W.).

## Author Contributions

C.B., K.L. and J.W. conceived the study and designed the experiments. C.B. performed the experiments. C.B., K.L. and J.W. wrote the manuscript.

## Data and materials availability

All data are available in the main text or the supplementary materials.

## Supplementary Figures

**Fig. S1:**
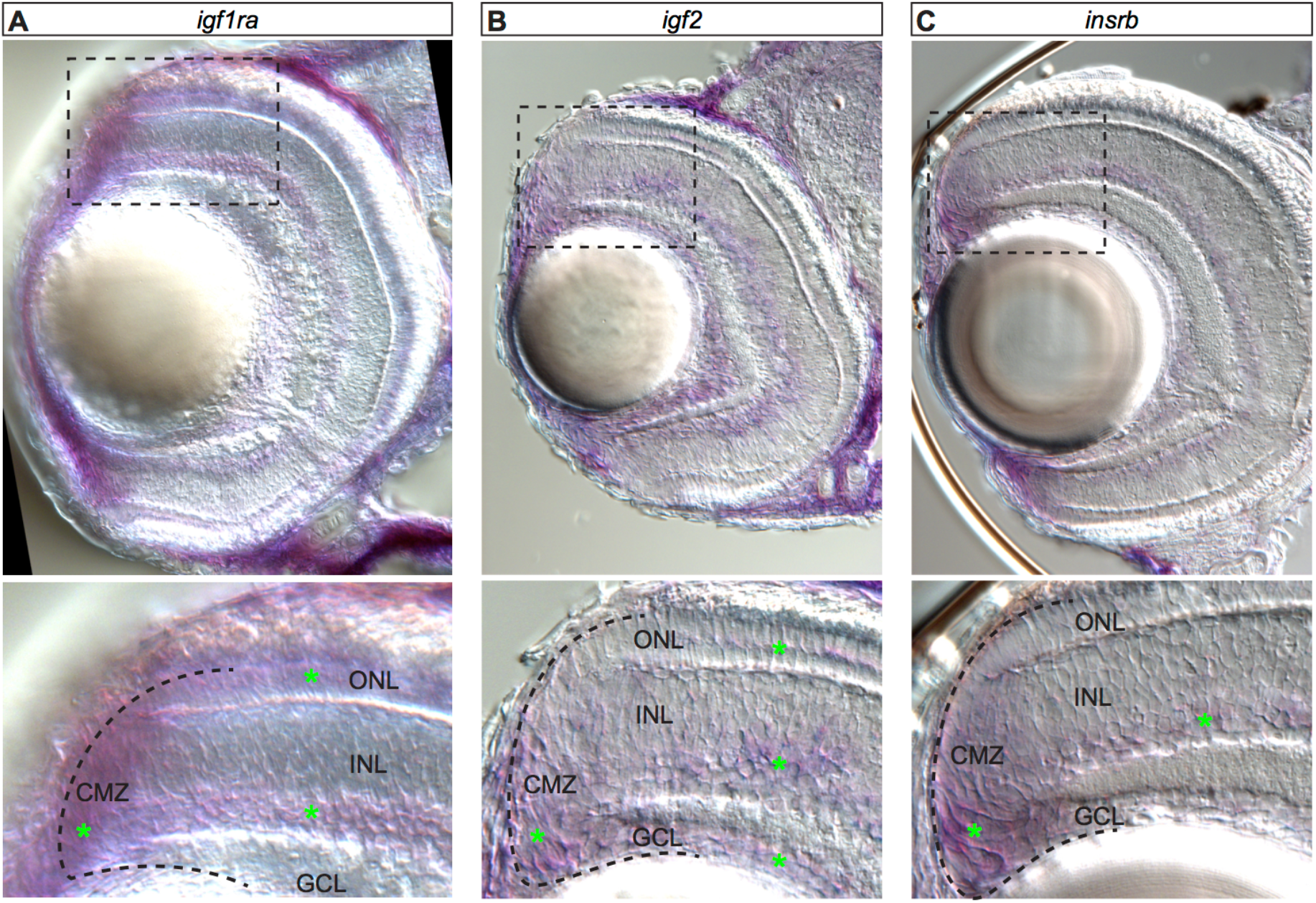
Igf signalling pathway components are expressed in the hatchling CMZ. (A-C) Cryosections of whole-mount *in situ* hybridisations of hatchling retinae. Expression of *igf1ra* (A) is visible in CMZ, outer nuclear layer (ONL) and INL (asterisks). *Igf2* (B) is expressed in CMZ, ONL, INL and ganglion cell layer (GCL). CMZ and INL show expression of *insrb* (C).

**Fig. S2:**
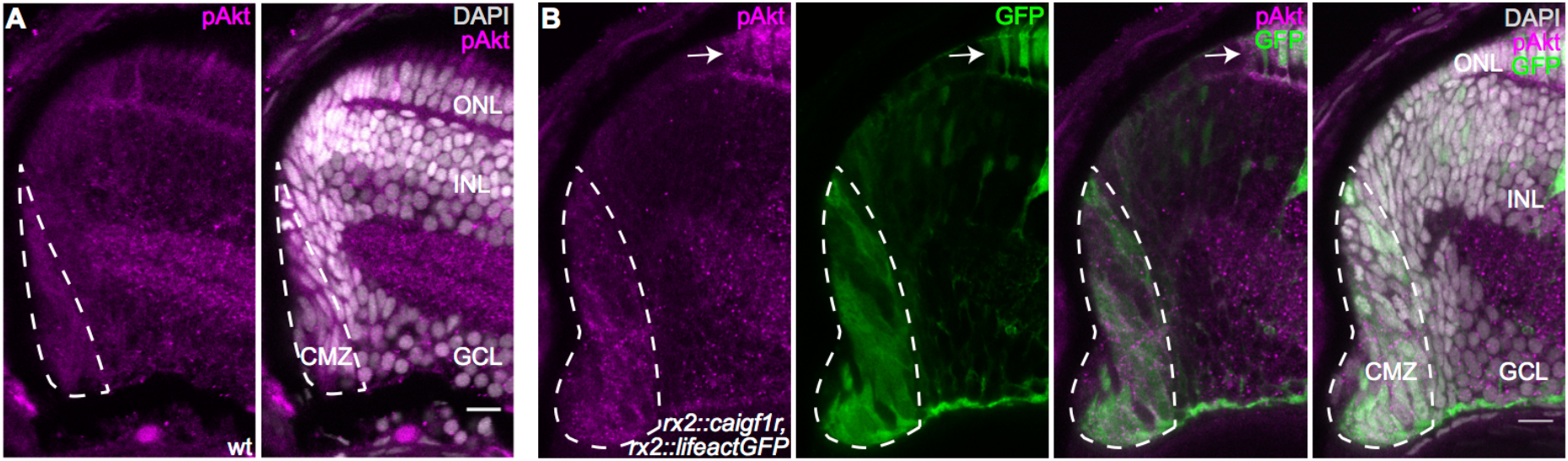
*Caigf1r* expression results in increased downstream signalling activation in the CMZ. (A-B) Cryosections of wt (A) and *rx2::caigf1r* (B) retinae at hatching stage. The pAkt-positive domain (magenta, dashed lines) is enlarged in *rx2::caigf1r* (B) compared to wt (A) retinae, co-localising with GFP signal (green) also in photoreceptors (B, arrow) (n = 3 fish each). Scale bars are 10 μm.

**Fig. S3:**
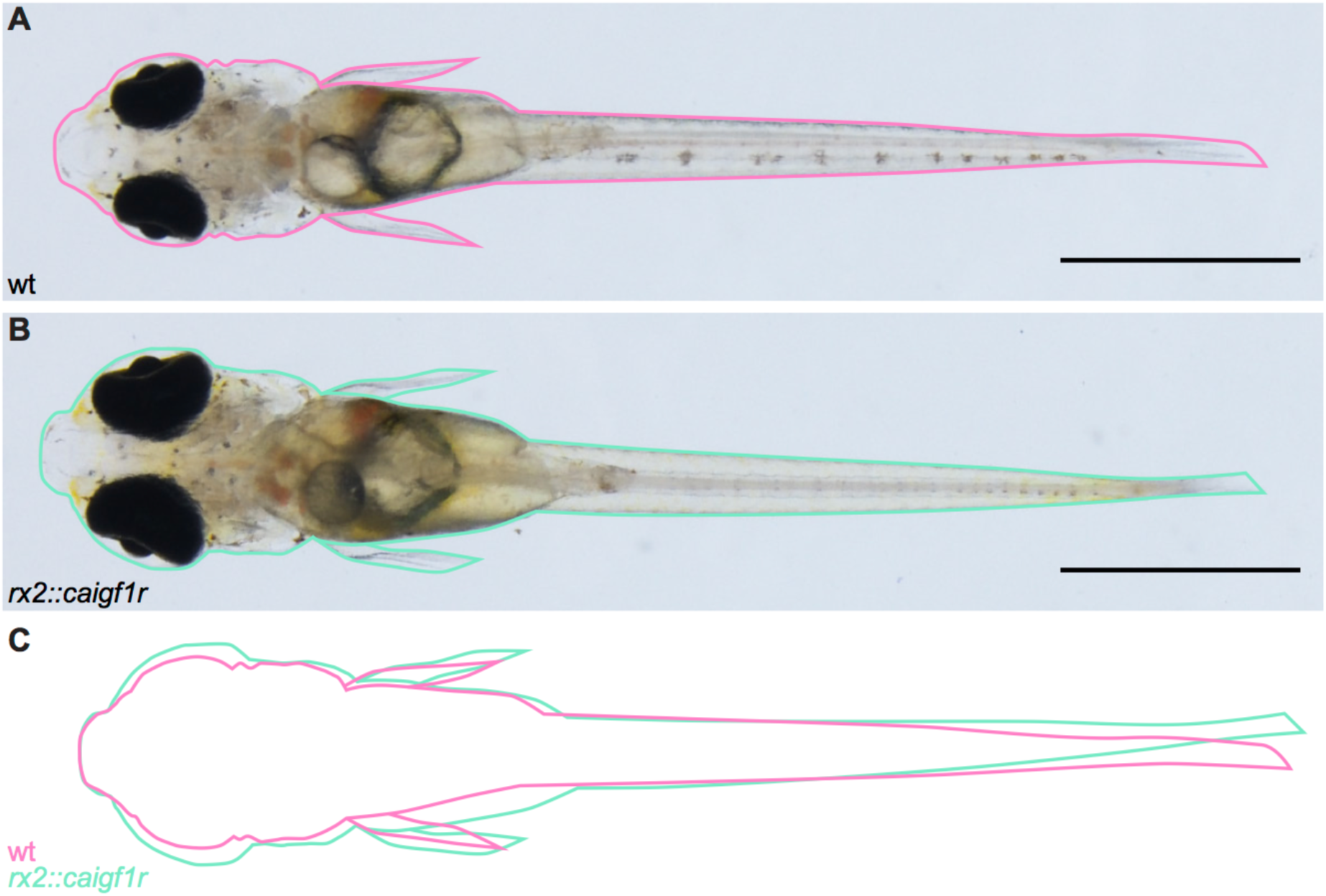
*Rx2::caigf1r* hatchlings are of comparable size to wt hatchlings. (A-B) Body size of *rx2::caigf1r* hatchling (B) is similar compared to wt sibling (A). (C) Overlay of body outlines of wt (A) and *rx2::caigf1r* hatchlings (B). Scale bars are 1 mm.

**Fig. S4:**
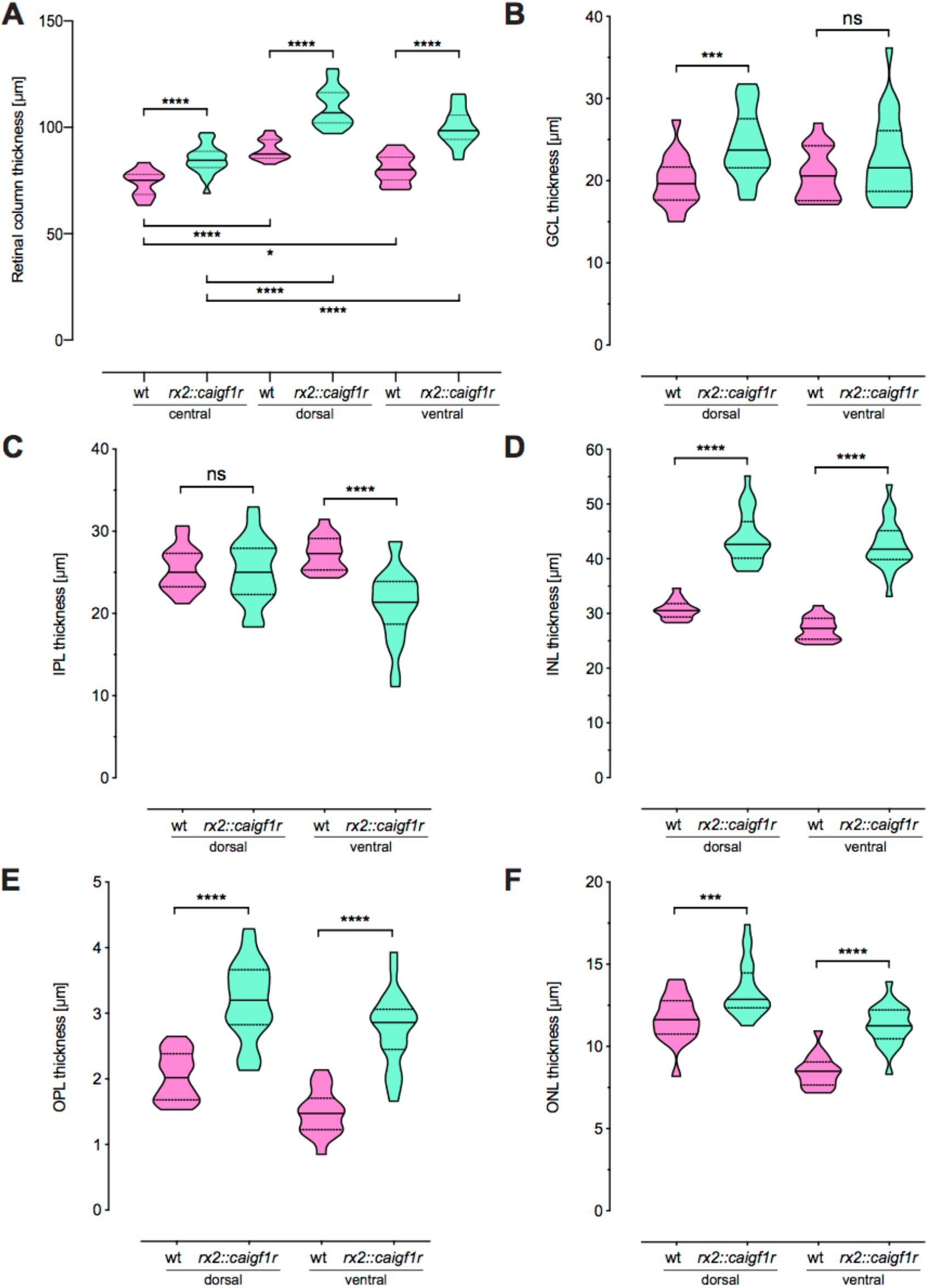
Neuroretinal thickness is increased throughout all nuclear layers. (A) Quantification of retinal column thickness in the central (wt: n = 11 sections from 8 retinae, *rx2::caigf1r:* n = 23 sections from 10 retinae), dorsal and ventral (wt: n = 18 sections from 12 retinae, *rx2::caigf1r:* n = 24 sections from 14 retinae) retina shows increase in *rx2::caigf1r* compared to wt fish and embryonic to CMZ-derived retina (median + quartiles, ****P < 0.0001, *P_wt c-v_ = 0.0130). (B-F) Quantification of dorsal and ventral individual retinal layer thickness in *rx2::caigf1r* (n = 24 sections from 14 retinae) compared to wt (n = 18 sections from 12 retinae) retinae (median + quartiles). Expansion of GCL in (B) (**P_d_ = 0.0046, ^ns^P_v_ = 0.1744), INL in (D) (****P_d/v_ < 0.0001), OPL in (E) (***P_d_ = 0.0002, ****P_v_ < 0.0001) and ONL in (F) (*P_d_ = 0.0427, ****P_v_ < 0.0001), but not IPL in (C) (^ns^P_d_ = 0.7863, ****P_v_ < 0.0001) is evident.

**Fig. S5:**
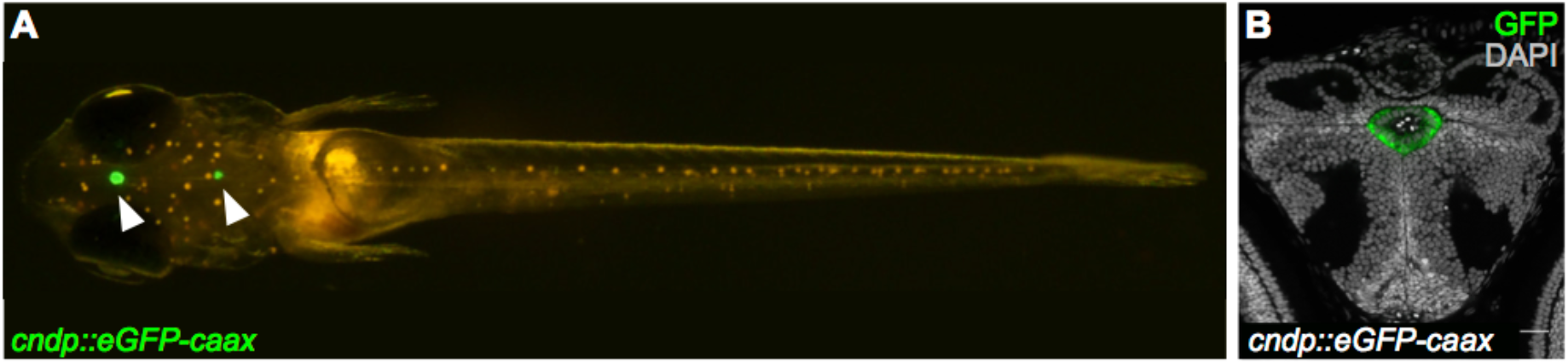
*Cndp* is expressed in the choroid plexi. (A) *Cndp::eGFP-caax* hatchling shows GFP expression in the choroid plexi in the brain (arrowheads). (C) Cryosection of a *cndp::eGFP-caax* hatchling brain. The diencephalic choroid plexus is positive for GFP (green). Scale bar is 20 μm.

**Fig. S6:**
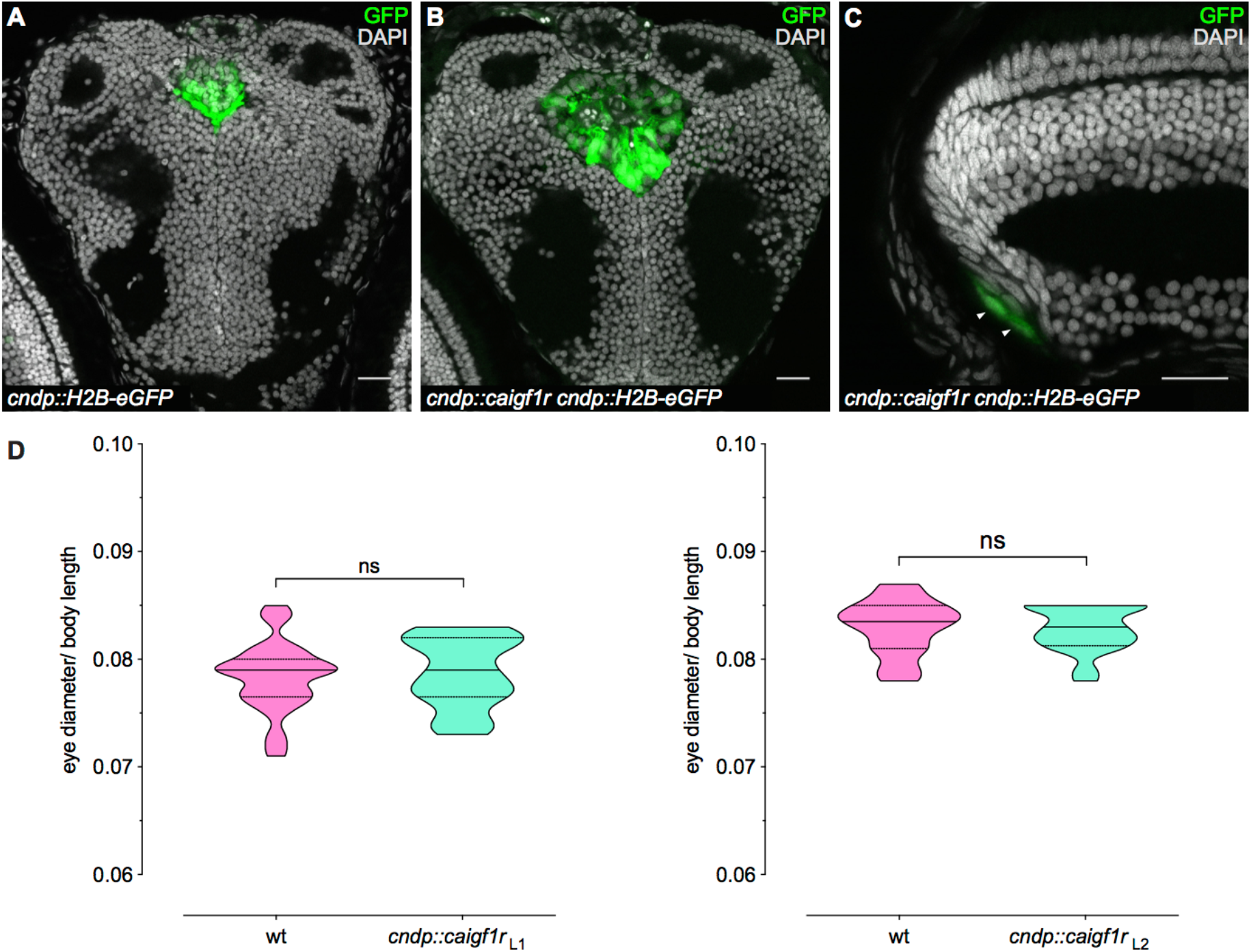
Expression of *caigf1r* in retinal stem cells does not result in increased eye size. (A-C) Cryosections of wt (A) and *cndp::caigf1r* (B-C) *cndp::H2B-eGFP* reporter hatchlings. The choroid plexi are positive for nuclear GFP (green) and enlarged in *cndp::caigf1r* (B) compared to wt (A) brains. *Cndp-* driven GFP expression (green, arrowheads) in the CMZ of *cndp::caigf1r* hatchlings (C) is not expanded (n = 3 fish). Scale bars are 20 μm. (D) Quantification of relative eye size (eye diameter normalised to body length) of wt (Line1: n = 29; Line2: n = 28) and *cndp::caigf1r* (L1: n = 17; L2: n = 20) hatchlings derived from two different founders (median + quartiles, ^ns^P_L1_ = 0.7457, ^ns^P_L2_ = 0.7590).

